# Sex inclusion in transcriptome studies of daily rhythms

**DOI:** 10.1101/2022.08.18.504312

**Authors:** Dora Obodo, Elliot H. Outland, Jacob J. Hughey

## Abstract

Biomedical research on mammals has traditionally neglected females, raising the concern that some scientific findings may generalize poorly to half the population. Although this lack of sex inclusion has been broadly documented, its extent within circadian genomics remains undescribed. To address this gap, we examined sex inclusion practices in a comprehensive collection of publicly available transcriptome studies on daily rhythms. Among 148 studies having samples from mammals in vivo, we found strong underrepresentation of females across organisms and tissues. Overall, only 23 of 123 studies in mice, 0 of 10 studies in rats, and 9 of 15 studies in humans included samples from females. In addition, studies having samples from both sexes tended to have more samples from males than from females. These trends appear to have changed little over time, including since 2016, when the US NIH began requiring investigators to consider sex as a biological variable. Our findings highlight an opportunity to dramatically improve representation of females in circadian research and to explore sex differences in daily rhythms at the genome level.

## Main Text

Underrepresentation of females is a persistent problem in biomedical research, and is particularly stark among preclinical studies using non-human mammals (Beery and Zucker, 2011; Woitowich *et al*., 2020). In addition, many preclinical studies either do not report the sex of biological samples or do not report results by sex (Mamlouk *et al*., 2020). Altogether, this lack of sex inclusion obscures the extent to which research findings generalize to roughly half the population (Clayton, 2016). Recognizing this issue, the US NIH created a policy in 2016 that requires investigators to factor biological sex into the design, analysis, and reporting of vertebrate animal and human studies (NOT-OD-15-102: Consideration of Sex as a Biological Variable in NIH-funded Research).

Unfortunately, lack of sex inclusion is an issue in circadian research as well. For example, only 34% of studies on the non-visual effects of light in humans have included females (Spitschan *et al*., 2022). Even more striking, among studies on light and circadian phase-shifting in rodents published 1964-2017, only 7% included females (Lee *et al*., 2021). The severe underrepresentation of female rodents in circadian research may be a legacy of early observations of the effects of the estrous cycle and estradiol on daily rhythms in hamsters (Morin *et al*., 1977; Takahashi and Menaker, 1980), although recent work indicates that female rats are not more variable than male rats in any neuroscience-related traits (Becker *et al*., 2016). Encouragingly, there is growing recognition of the importance of addressing sex bias and investigating sex differences in sleep and circadian rhythms (Spitschan *et al*., 2022; Joye and Evans, 2022). Although sleep and circadian research increasingly makes use of genomic techniques, sex inclusion trends in genomic studies of daily rhythms remain undescribed.

To address this gap, we first assembled a collection of publicly available transcriptome studies (bulk microarray or RNA-seq) from mice, rats, or humans that had samples from at least three times of day (Tables S1 and S2). We downloaded the metadata for each study using the seeker R package (https://seeker.hugheylab.org). From the metadata, we extracted the organism, tissue, time of day, and biological sex (where available) of each sample from each study. Where necessary, we obtained information on biological sex from the published article. For our analysis, we only included studies linked to a published article, and considered one study as corresponding to one article. We did not filter studies or samples with respect to genotype, light-dark cycle, or any other experimental condition. Altogether, the collection comprised 123 studies in mice (7,305 samples), 10 in rats (373 samples), and 15 in humans (191 subjects). We defined each study’s sex inclusion status as male only, female only, male and female (if each sample’s sex was identifiable), mixed (if each sample was based on pooled tissue from males and females, or if the article stated using both males and females, but each sample’s sex was not identifiable), or unspecified.

We examined sex inclusion by organism (Fig. 1A and Table S3). In mice, 91 of 123 studies (74%) included only males, whereas 23 studies (19%) included females, either as female only, male and female, or mixed. In rats, all nine studies having specified sex included only males. In humans, sex inclusion was more balanced, as 8 of 15 studies had samples from males and females. We observed similar trends when analyzing the collection based on numbers of samples (Table S4). These results indicate that females are highly underrepresented in mammalian circadian transcriptome data.

**Fig. 1.**
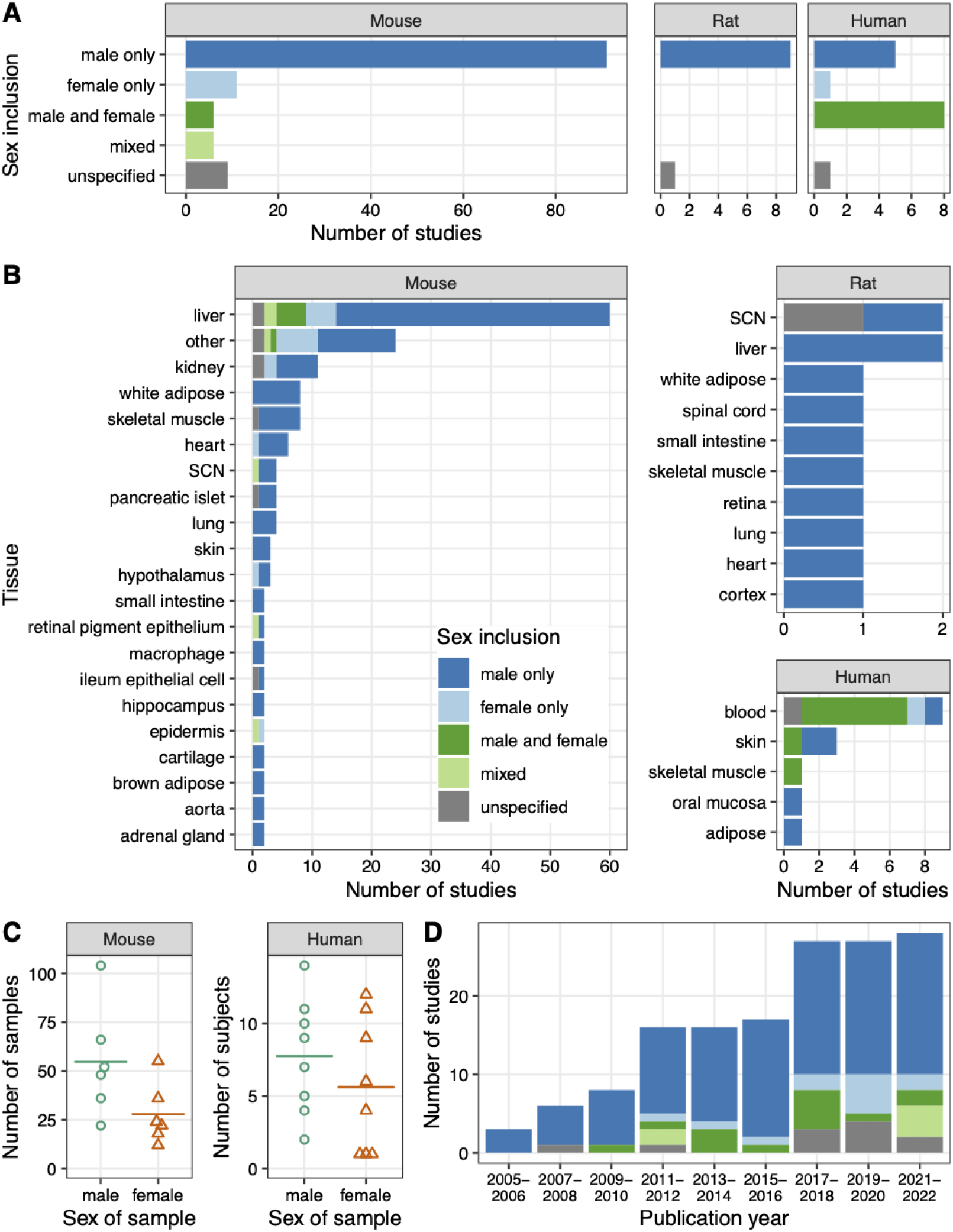
Sex inclusion in circadian transcriptome studies from mice, rats, and humans. Each study corresponds to a published article. **(A)** Barplots of number of studies by organism and sex inclusion status. The latter is also represented by color, which is consistent throughout the figure. **(B)** Barplots of number of studies by organism, tissue, and sex inclusion status. SCN indicates the suprachiasmatic nucleus. For mouse studies, the “other” tissue comprises tissues represented in only one study. Some studies included samples from multiple tissues, so the total counts in (B) could be more than the counts in (A). **(C)** Beeswarm plots of number of samples (per mouse study) and number of subjects (per human study) among studies whose sex inclusion status was male and female. Each point represents a study. Each horizontal line represents the mean for that group. **(D)** Barplot of number of studies (from all organisms) by sex inclusion status and publication year of the corresponding article.

We next examined sex inclusion by organism and tissue (Fig. 1B and Tables S5 and S6). In mice, 23 of 44 unique tissues had samples only from males, whereas 16 tissues had samples in some way from females (as female only, male and female, or mixed; Table S7). Within liver, by far the most common tissue, 46 of 60 studies were male only, whereas 5 were female only and 5 were male and female (Table S8). In humans, the most common tissue was blood, for which 6 of 9 studies included males and females. However, both mouse and human studies including males and females tended to have more samples (or subjects) from males than from females (Fig. 1C).

We also examined sex inclusion trends over time, based on the publication year of each study’s corresponding article (Fig. 1D and Table S9). These results indicate that the underrepresentation of females has remained roughly constant since 2011, although the number of studies having “mixed” sex inclusion appears higher since 2021.

To explore these results further, we examined the articles for studies published in 2021-2022, which revealed three main findings (Table S10). First, most studies whose sex inclusion status was male only or female only did not mention a justification. Second, the underrepresentation of females in transcriptome experiments was concordant with that of other experiments. Third, of the nine studies that included males and females for any experiments, two reported results by sex.

Although the collection of studies we examined here is extensive, it is not exhaustive. First, it includes only transcriptome studies, not studies based on other genomic techniques, which are relatively less common and often performed on the same or similar samples. Second, the collection only includes studies in which time of day was an experimental variable, and thus excludes studies based on tissue from postmortem human donors, where information on time of day of death has revealed daily variation in gene expression and where donors may be more sex-balanced (Li *et al*., 2013; Chen *et al*., 2016; Ruben *et al*., 2018). Third, the collection does not include studies from less commonly used vertebrates or from invertebrates—despite the importance of fruit flies to circadian research. Nonetheless, our findings are consistent with those for biomedical research as a whole (Woitowich *et al*., 2020) and for circadian phase-shifting experiments (Lee *et al*., 2021).

Our study highlights the utility of compiling and standardizing the vast amount of publicly available circadian data (Pizarro *et al*., 2013; Ceglia *et al*., 2018), as also recently shown by a meta-analysis of circadian gene expression in mouse liver (Brooks *et al*., 2022). Indeed, a secondary finding of our analysis is the predominance of liver as a tissue source among mouse studies. Although the liver may have somewhat more rhythmic genes compared to other tissues (Zhang *et al*., 2014), its current popularity is likely out of proportion to its importance in the mammalian circadian system, even among peripheral tissues. A similar argument applies to human blood, although here the primary driver is likely the relative non-invasiveness of drawing blood from live humans.

Given the few studies and tissues having sufficient data from males and females, we leave a meta-analysis of sex differences in daily rhythms for future work. Previous work indicates that some differences may be subtle, others less so (Weger *et al*., 2019; Joye and Evans, 2022). To this point, a sex-based analysis is not possible for studies whose sex inclusion is “mixed” (whether due to pooling or lack of labeling). Rigorous quantification of sex differences in circadian genomic data will entail not only proper experimental design, but also use of statistically valid methods for quantifying differential rhythmicity, in order to avoid misinterpretation (Thaben and Westermark, 2016; Singer and Hughey, 2018; Weger *et al*., 2021; Pelikan *et al*., 2021). Importantly, however, the ethical principle of fair representation—as well as the NIH’s policy on sex as a biological variable—does not require that all studies be powered to detect sex differences. Such a requirement could be challenging for circadian genomic experiments, which already involve a time-course, potentially in multiple conditions, on limited budgets. Instead, the principle and policy require that sex be factored into every step of research.

In summary, females remain strongly underrepresented in circadian genomic studies. Given the duration of funding cycles and peer review, one might not expect the 2016 NIH policy to have an immediate effect. However, the apparent lack of justification for decisions on sex inclusion, even in recent studies, points to an opportunity for us as a research community to do better. This opportunity seems especially relevant as the field moves increasingly from bulk to single-cell genomics. Improving sex inclusion could both improve our work’s generalizability and contribute to a growing understanding of the role of biological sex in daily rhythms.

## Acknowledgments

This work was supported by the U.S. National Institutes of Health R35GM124685 (to JJH) and F31GM143909 (to DO).

## Data availability

Reproducible results are available at https://doi.org/10.6084/m9.figshare.20502372.

